# Hippocampal inactivation during rearing on hind legs impairs spatial memory

**DOI:** 10.1101/2022.10.21.513207

**Authors:** D.M. Layfied, N. Sidell, K. Blankenberger, E.L. Newman

## Abstract

Spatial memory requires an intact hippocampus. Hippocampal function during epochs of locomotion and quiet rest (e.g., grooming and reward consumption) have been the target of extensive study. However, during navigation rats frequently rear up on to their hind legs and the importance of hippocampal activity during these periods of attentive sampling for spatial memory is unknown. To address this, we tested the necessity of dorsal hippocampal activity during rearing epochs in the study phase of a delayed win-shift task for memory performance in the subsequent test phase. Hippocampal activity was manipulated with closed-loop, bilateral, optogenetic inactivation. Spatial memory accuracy was significantly and selectively reduced when the dorsal hippocampus was inactivated during rearing epochs at encoding. These data show that hippocampal activity during periods of rearing can be important for spatial memory, revealing a novel link between hippocampal function during epochs of rearing and spatial memory.

## Introduction

Spatial memory is a core cognitive ability that is central to many daily activities. Hippocampal integrity has long been recognized to be essential for spatial memory^1–7^. Yet, not all epochs of hippocampal processing are equally important for spatial memory encoding. Observations of spatial tuning during locomotion and of replay during periods of quiet rest (e.g., grooming and reward consumption) have made these behavioral epochs the focus of most work studying how hippocampal processing supports spatial memory^8–13^. Yet, animals of all types perform behaviors wherein they stop locomotion and, distinct from quiet rest, actively sample the surrounding environment^14^. In mammals, including rats and mice, this can appear as rearing onto their hind legs^15^. The relevance of rearing for hippocampal dependent encoding of spatial memories is unknown.

Rearing, defined as when an animal stands up onto its hind legs, is a widely observed but minimally researched behavior. Rearing offers an animal a distinct perspective for sensory sampling (visual, olfactory, etc.), putatively offering improved information about distal cues to support environmental modeling and rearing reliably increases in response to environmental novelty^15–19^. Lever and colleagues^15^ synthesized what little was known in this regard to hypothesize that rearing is an ethological measure that could be used to assess hippocampal learning and memory. However, it remains untested whether hippocampal activity during rearing substantially contributes to spatial memory encoding.

What little is known of hippocampal function during rearing supports the hypothesis that rearing is an epoch of hippocampal encoding. Functional recordings of the dorsal hippocampus during rearing show that it is associated with increased power of 7-12 Hz ‘high theta’ rhythmic activity^20–24^. This supports the hypothesis that rearing is an epoch of hippocampal processing because theta, itself, is associated functionally with hippocampal encoding and retrieval^25–34^, disrupting theta interferes with hippocampal dependent memory^35–38^, and restoring theta can rescue hippocampal dependent learning deficits^39,40^. Yet, studies examining theta and its relationship to learning and memory are standardly restricted to periods of normal horizontal locomotion during spatial exploration. Thus, while the correlation between hippocampal theta and rearing suggests that rearing could be an epoch of hippocampal dependent encoding, the relevance of rearing for spatial memory is unknown.

These convergent reasons led us to hypothesize that rearing is an epoch of hippocampal dependent encoding of spatial memories. To test this hypothesis, we selectivity inactivated the dorsal hippocampus during rearing events in a spatial memory task. Hippocampal activity was manipulated with closed-loop, bilateral, optogenetic inactivation, triggered by a 3D camera system calibrated to detect rearing. Manipulations of hippocampal activity were restricted to the study phase of an 8-arm maze delayed-win-shift task^41,42^. Behavioral assessments were performed during the subsequent test phase. We found that inactivation of the dorsal hippocampus during bouts of rearing resulted in impaired spatial memory at test. When the hippocampus was inactivated for matched amounts of time but at a delay relative to rearing events, we did not find significant memory impairments. This effect was unique to the experimental rats that were transfected to express halorhodopsin and a fluorescent reporter. No impairments were observed in any condition in control rats that were transfected to express the fluorescent reporter only. These data provide the first evidence that the activity of the dorsal hippocampus during rearing is important for spatial memory.

## Methods

### Animals

All animal procedures and surgeries were conducted in strict accordance with National Institutes of Health and the Indiana University Institutional Animal Care and Use Committee guidelines. Adult (at least 3 months of age) Long Evans rats were used for all experiments. A total of 13 rats were used: 6 male rats were in the experimental group and 7 rats (4 female; 3 male) were in the control group. Animals were individually housed and maintained on a 12 H light/dark cycle in a temperature and humidity-controlled room with ad libitum access to water, and food restricted to maintain ~90% (85-95%) of free feeding body weight. Rats were acclimated to the animal facility for 5 days before being handled daily.

### Apparatus

Training took place on a custom built automated 8-arm radial maze. The maze had a 33.2 cm wide hub and white opaque pneumatic drop doors at the entrances to the 8 arms. The arms measured 48.26 cm long, 10.79 cm wide and had opaque white acrylic floors. The arms had 20.95 cm tall clear acrylic walls allowing for viewing of distal cues (wall decorations, bookshelf, desk, door) surrounding the maze at a range of distances from one to five feet from the maze. At the end of each arm were food wells in which 45 mg sucrose pellets (Bio-Serv, Flemington, NJ) were delivered. The maze was open on top to permit testing of tethered animals. The maze was cleaned with chlorhexidine immediately after each trial.

### Experimental design

The experimental design was a 2×3 design with two levels for opsin expression {opsin + reporter, reporter only} manipulated between animals and three levels for light delivery timing {’Off’, ‘Rear’, ‘Delay’} manipulated within animal. Manipulations of light delivery timing served to test the relevance of hippocampal activity during rearing behavior. ‘Off’ established the baseline ability of each rat to complete the task. ‘Rear’ measured the impact of timing light delivery to be synchronous with rearing and represented the key experimental condition. ‘Delay’ controlled for the total amount of light delivered within a trial but desynchronized the light delivery from the rearing behavior. Each rat completed 6-14 trials of each optogenetic condition in pseudo random order. The only constraint on the order was that all three conditions be tested before any was repeated another time. Manipulations of opsin expression served to isolate the effect of the opsin on behavior by controlling for any incidental effects of light delivery or viral transfection. The opsin expression conditions formed two cohorts of rats, one of which was transfected with the opsin and fluorescent reporter while the other was transfected with reporter only.

### Behavioral training

#### Habituation & preliminary training

For 3 days prior to training, rats were handled for 10 min and given 20–30 sucrose pellets to habituate them to experimenter handing and rewards. Preliminary training consisted of one 10 min. trial daily for 4 days. During each, 4–6 pellets were placed along each of the 8 arms and 2 pellets in the food wells of each arm. The rat was allowed to freely explore the full maze. The trial ended after the rat consumed all pellets or 10 min had elapsed.

#### Initial training

Initial training consisted of 10 sessions, one trial/session. During each, rats were trained that arms would be baited once per day. Prior to each trial, the food wells of all 8 arms were baited with 2 pellets each. Training trials began by placing the animal in the central hub with hub doors closed. After one minute, the doors opened allowing the rat to freely forage. The rats could explore until all pellets were collected or 15 min. had elapsed.

#### Task

The spatial memory task used here was the delayed-win-shift task on an 8 arm radial arm maze. The task consisted of three phases, a study phase, a delay phase, and a test phase as shown in Figure 1. In the study phase, the rat was placed in the central hub. After 30 seconds a random set of four doors opened and the rat was allowed to collect pellets from each. The rat was then removed from the maze and placed on pedestal next to the maze for a 4 min. delay phase. During which, the maze was cleaned with chlorhexidine to remove odor cues. The test phase began by placing the rat in the hub and. After 30 seconds, all eight doors opened. The four arms that had not opened during the study phase baited food wells. The test phase ended after all the pellets had been consumed or when 15 min elapsed. Regular training (~5 days/week) continued until rats achieved behavioral criterion: no more than three errors over four days.

**Fig. 1.**
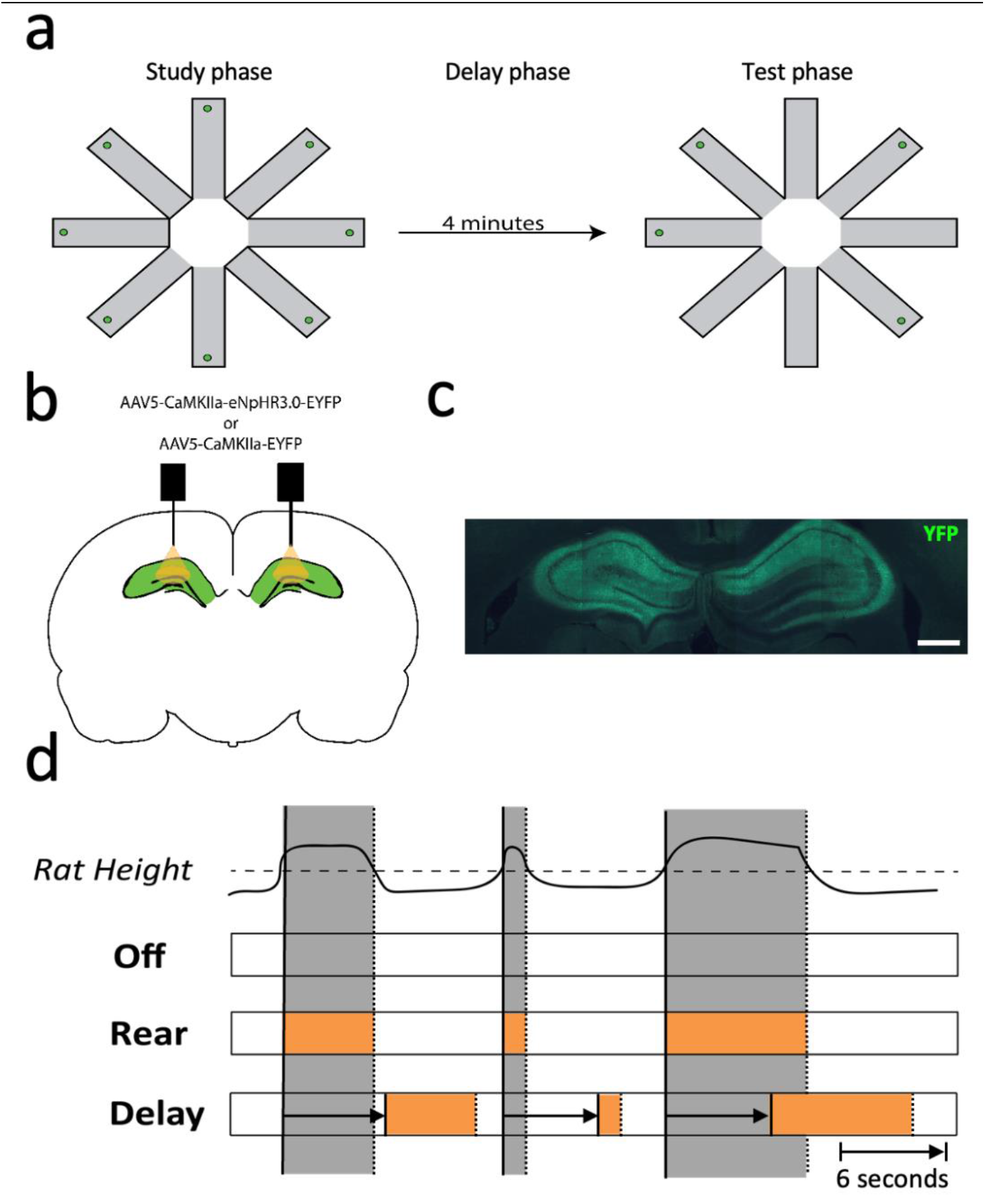
Overview of the experimental paradigm. **a** A delayed-win-shift task was used. In the study phase, 4 doors opened and the rats foraged for rewards in the open arms. In the delay phase, the rat was removed from the maze for 4 min. and the maze was cleaned. In the test phase, all 8 doors opened and rats foraged for the remaining rewards. **b** Dorsal hippocampus was targeted with viral infusions and optical fiber implants **c** Immunofluorescence labeling EYFP expression induced in the dorsal hippocampus. Scale bar is 1 mm. **d** Three optogenetic stimulation conditions were used. Top: Black line plots a hypothetical trace of a rat’s height over time. Grey zones mark rearing events, when the rat’s height crosses a fixed threshold (dashed line). Off: A control condition wherein no optogenetic stimulation occurred. Rear: The laser is activated (orange rectangle) during rearing events. Delay: The laser is activated based on rearing but the onset and offset occur with a fixed 6 second delay as indicated by the horizontal arrow.

### Behavioral scoring

The key dependent measures of spatial memory performance were percent correct and number of arm entries. Percent correct was measured as the number of the first four choices of the test phase that contained rewards divided by four. Number of arm entries was measured as the total number of arms entered to find all four baited reward sites. A visit to an arm was defined to occur when the rat’s hind feet entered the arm. An arm visit counted as an error if the arm had been visited earlier in the trial.

### Rear detection

Rearing behavior was tracked using a 3D camera (RealSense Depth Camera D435; Intel). The camera was positioned on the ceiling above the center of the maze facing down such that it was able to capture the whole of the maze at 30 Hz and 640 × 480 resolution. A custom real-time analysis package was written to detect rearing events. Rearing events were defined as moments when a ‘blob’ of appropriate size entered a region of interest. The region of interest was a 3D zone filling a short and broad cylinder of space above the maze. The cylinder perimeter matched the perimeter of the maze enclosure. The bottom of the cylinder was aligned with the top of the enclosure walls. The top of the cylinder extended 20 cm above this. The top limit was defined to prevent erroneous triggering events from the tether as it moved between the camera and the enclosure. To further prevent erroneous triggering, blobs were required to occupy more than 0.13 cm^2^ and less than 26.46 cm^2^. This served to prevent the experimenter from incidentally triggering the laser by coming into view of the camera. The rear detection routine was implemented with the Open-source Computer vision’s (OpenCV) blob detector. When a blob of appropriate size was detected in the region of interest, a command is sent to an Arduino unit which then activated the laser system. For control rats the 3D camera laser system was operated manually.

### Optogenetic control and light delivery

Optogenetic control was implemented with the eNpHR3.0 halorhodopsin, a light gated ion pump which inhibits neural activity with photostimulation^43^. Activation of halorhodopsin was achieved using light from a CE:YAG laser diode optical head laser system (Doric) filtered to 570 nm to 615 nm. Laser output was delivered to the hippocampus by way of a patch fiber connected to a rotary joint (Doric) and, from the rotary joint a dual fiber optic patch cord (Doric) that was coupled to the implanted optical fibers prior to each testing session. Light intensity was controlled by the Doric Neuroscience studio software to obtain 5-10 mW at the tip of the fiber in the brain using a photodiode power sensor coupled to a power meter (Thorlabs). Laser activation was triggered by an external control signal generated by an Arduino unit (Arduino due) that was under the control of the custom rearing detection software.

The optogenetic experimental conditions were as follows: In the main experimental condition wherein light was delivered coincidentally with rearing behavior, referred to as ‘Rear’ in the results, the laser was activated as the 3D camera system detected rearing and remained on for the full duration of the rear. In the baseline ‘Off’ condition, the main power switch for the laser remained off so that no light was delivered at any point. Nonetheless, the optical fiber was attached as in the other conditions. In the control ‘Delay’ condition, light was delivered in response to rearing but both the laser activation and deactivation were triggered at a fixed 6 second delay relative to the rearing. The 6 second delay was inserted by the control software between the time when the 3D camera system detected rearing and time when the state of laser system was switched. Thus, importantly, the duration of light delivery in the ‘Delay’ condition matched the duration of the detected rear. All rats were coupled to the fiber patch cords during all testing, irrespective of condition or cohort.

### Stereotaxic surgery and viral vectors

Rats underwent two surgeries. In the first, a virus was injected into the dorsal hippocampus. In the second, an optical fiber cannula for light delivery and opsin activation was implanted. During the first surgery, rats were anesthetized with 1.5-4% isoflurane and their head was positioned in a stereotaxic frame. The scalp was cut and retracted. Three sites were drilled above the hippocampus bilaterally. In each, viral infusions were performed at 3 different depths. Thus, a total of 18 separate 45 nl injections were done at the following coordinates: [-3.0 AP, +/-2.2 ML, 2.1, 2.3, 2.5 DV]; [-3.7 AP, +/-2.9 ML, 2.0, 2.2, 2.4 DV]; and [-4.3 AP, +/3.5 ML, 2.0, 2.2, 2.4 DV]. Halorhodopsin and fluorescent reporter expression were transduced with AAV(5)-CaMKIIa-eNpHR3.0-EYFP (UNC vector core). The same AAV serotype and promoter have been used to induce opsin expression in the rat hippocampus previously^44^. At the end of surgery, the scalp was sutured shut and the rat was allowed to recover for one week post surgery before continuing regular behavioral training. Rats underwent the second surgery 2-5 weeks after the first surgery. Again, rats were anesthetized with 1.5-4% isoflurane, their head was mounted in a stereotaxic frame, and the scalp was cut and retracted. Two optical fibers (MFC_200/245-0.53_5mm_MF2.5-FLT; Doric Inc) were positioned above the center injection sites from the previous surgery, at -3.7 AP, 2.9 ML, 1.8 DV, and fixed to two jewelers screws inserted into the skull with dental acrylic. The scalp was sutured closed around the implant. Control rats underwent one combined viral injection and optical fiber implantation surgery. Only a fluorescent reported was expressed by infusing the virus AAV(5)-CAMKIIa-EYFP (UNC vector core). One 2 μl injection into each side of the hippocampus [-3.6 AP, +/- 2.8 ML, 2.4 DV] with two optical fibers (MFC_200/245- 0.53_5mm_MF2.5-FLT; Doric Inc) placed bilaterally above the injection site. Again, rats were allowed to recover for one week following surgery before restarting behavioral training. Data collection began after rats reached behavioral criterion.

### Histology and immunohistochemistry

Upon completion of testing, animals were euthanized via isoflurane overdose and perfused intracardially with phosphate buffered saline (PBS) followed by a 4% paraformaldehyde saline solution. Brains were saturated with a 30% sucrose solution prior to sectioning. Coronal sections (50 um thick) were cut with a crysostat (Leica Biosystems) or microtome (American Optic company). Immunohistochemistry was performed on free-floating sections to amplify induced EYFP reporter signaling. Sections were first rinsed with PBS then blocked with buffer (PBS, 5% normal goat serum, and 0.4% Trition X-100). This was followed by overnight incubation with conjugated anti-GFP rabbit antibody (1:1000; catalog no. A21311; Invitrogen). Finally, the sections were rinsed with PBS and then mounted on slides and cover slipped with DAPI and Fluoroshield.

### Statistical analysis

Because individual animals completed different numbers of testing sessions, statistical analyses were performed with a multilevel hierarchical model to test the relationship between the light off condition and optogenetic stimulation change in measures of interest. The model treated light delivery timing as a fixed effect and rat identity was set as a random effect. Concretely, the model was implemented with the equation “*Perf* ~ *Opto* + (*Opto|Rat*)” where *Perf* is the behavioral score of interest (percent correct or number of arms entered), *Opto* was a categorical variable indicating the light delivery timing condition and *Rat* was an index indicating the rat identity from which each data point came. Models were analyzed in MATLAB using the *glme.m* function. We sought to determine whether the slope relating to *Opto* to *Perf* was significant that would indicate that the timing of light delivery changed the respective behavioral score. Results are reported as the expected effects and 95% confidence intervals (e.g., Effect [lower bound, upper bound]) as determined by the hierarchical modeling. Significance was determined based on an alpha level of 0.05.

## Results

To determine if hippocampal activity during rearing is important for spatial memory, we performed a closed-loop experiment wherein we tested the effect of optogenetically inhibiting the dorsal hippocampus during epochs of rearing on spatial memory. Spatial memory was assessed with the eight-arm delayed-win-shift task, consisting of study, delay and test phases (Fig. 1a). During the study phase, four of eight arms were opened to the rat and, in each, the rat found food rewards. In the delay phase, rats were removed from the maze for four minutes. Finally, in the test phase, rats were granted access to all eight arms but could only find rewards in the four previously unopened arms. Rats were trained on this task to criterion level accuracy (<3 errors in four consecutive days of testing, >80% accuracy) before and after the viral transfection and fiber cannula implantation surgeries. Viral transfection targeted the bilateral dorsal hippocampus proper and either transduced expression of halorhodopsin and fluorescent reporter (experimental group) or fluorescent reporter only (control group). Fiber optic cannula were implanted bilaterally dorsal to the viral injection (Fig. 1b & 1c). Epochs of optogenetic manipulation were restricted to occur in the study phase only. Hippocampal activity was unmanipulated during the test phase when spatial memory was assessed (Fig. 1d).

Starting with the experimental group, comparing percent correct at test between the ‘Rear’ condition, wherein light delivery was synchronized to rearing, and the ‘Off’ condition, wherein no light was delivered, revealed a significant reduction to spatial memory (77.7% vs. 65.7%; GLME est. = -11.9% [-21.6%, -2.3%], *t(178) = -2.44, p = 0.02).* The same comparison performed on data collected from the control group, wherein halorhodopsin was not expressed, revealed that light delivery had no significant effect on percent correct (81.4% vs. 83%; GLME est. = 2.7 [-4.1, 6.5], *t(217) = 0.4, p = 0.66).* These data are shown in Fig. 2a.

**Fig. 2.**
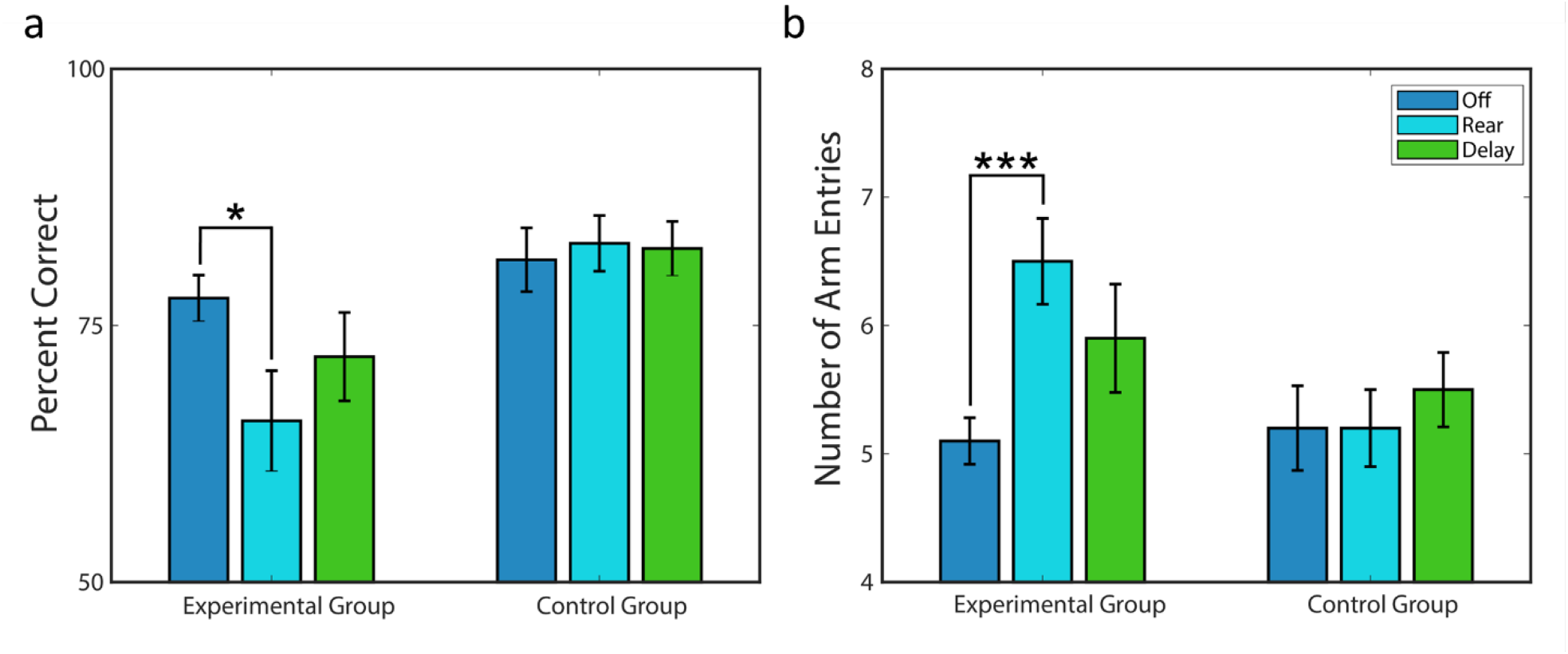
Optogenetic inactivation of dorsal hippocampus during rearing impairs spatial memory. **a** Effect of optogenetic inhibition on the percent correct measure across experimental and control groups **b** Effect of optogenetic inhibition on number of arm entries during test across experimental and control groups. *P<0.05; ***P<0.001; mean ± SEM *n* = 6, experimental group; *n* = 6, control group.

Examining the number of arms entered to find all rewards instead of percent correct revealed the same pattern of results. For the experimental group, rats entered significantly more arms while finding all the rewards on ‘Rear’ trials than ‘Off’ trials (5.1 vs. 6.5; GLME est. = 1.4 [0.78, 2.1], *t(178) = 4.3, p < 0.0001).* Rats in the control condition were not significantly affected by the light delivery in the ‘Rear’ trials relative to the ‘Off’ trials (5.2 vs. 5.2; GLME est. = 0.002 [-0.59, 0.59], *t(217) = 0.009, p = 0.99*). These data are shown in Fig. 2b.

Differences between the ‘Rear’ and ‘Off’ conditions could have been simply due to having intermittently disrupted hippocampal activity during the study phase of ‘Rear’ trials. That is, the ‘Rear’ versus ‘Off’ comparison does address whether it mattered that inactivation was synchronized to rearing. To test if synchronization of the light delivery to rearing was a determining factor in the effects listed above, we also tested whether inserting a six second delay between detected rearing events and the laser activation would also disrupt spatial memory performance. This is the ‘Delay’ condition. If the memory impairments observed above were simply the result of inactivating the hippocampus at all, irrespective of whether the rat was rearing, then we should expect a significant impairment in the ‘Delay’ condition as well. However, if the memory impairments resulted because we synchronized the inactivation with rearing behavior, then we would not expect a significant impairment when the laser activation is delayed relative to rearing.

The analyses showed that performance in the ‘Delay’ was not significantly different from the ‘Off’ condition with regard to percent correct (77.7% vs. 72.0%; GLME est. = -5.7 [-14.2, 2.8], *t(178) = -1.3, p = 0.19)* and only a trend toward an increase in total arm entries (5.1 vs 5.9; GLME est. = 0.8 [0.0, 1.7], *t(178) = 2.0, p = 0.05).* Note, this trend is in contrast to the strong effect observed when comparing the ‘Rear’ condition to the ‘Off’ condition. That the inactivation should show any trend is not surprising, but it is also likely that the six second delay did not fully desynchronize light delivery from rearing behavior. As expected, but included here for full transparency, the delay condition similarly had no significant effect in the control group for percent correct (81.4 vs 82.4%; GLME est. = 2.6 [-4.1, 6.3], *t*(217) = 0.41, *p* = 0.68) or total arms entered (5.2 vs. 5.5; GLME est. = 0.3 [-0.2, 0.92], *t*(217) = 1.17, *p* = 0.24). ‘Delay’ condition data are shown in Fig 2.

## Discussion

Here, we sought to test the relevance of rearing as an epoch of hippocampal dependent encoding of spatial memories. To address this, we examined how spatial memory was affected by closed loop optogenetic inhibition of the dorsal hippocampus activity during rearing. The results show that spatial memory performance in the test phase of a delayed radial maze win-shift task decreased significantly when dorsal hippocampus activity was inhibited selectively during rearing in the study phase. Spatial memory impairments were not observed when a control virus that lacked the halorhodopsin gene was used ruling out the possibility light delivery effects (e.g., tissue heating, visual distraction, etc.) caused the reduced performance in the experimental group. The fact that the ‘Delay’ condition also didn’t generate significant performance impairments indicates that it is not simply inactivating the hippocampus at odd intervals during the study phase that impaired memory. Together, these results demonstrate that dorsal hippocampal activity during rearing is important for spatial memory.

Our results are consistent with the hypotheses of Lever and colleagues regarding the functions of rearing. Having reviewed the limited body of work on rearing, Lever and colleagues^15^ synthesized the available data with the hypothesis that ‘*rearing is a useful marker of environmental novelty, that the hippocampal formation is a crucial component of the system controlling rearing in novel environments, and that rearing is one of several ethological measures that can profitably be used to assess hippocampal learning and memory*.’ Strong evidence was available at the time to support their hypothesis that rearing marks environmental novelty: A wide variety of mammals rear in response to environmental novelty. The hypothesized links to hippocampal function were more speculative. The data indicated that lesions of the hippocampus have inconsistent effects on rearing frequency^45,46^. Only indirect evidence existed to link rearing and hippocampal learning. For example, rearing covaried with performance in the Morris water maze—declining during learning and reinstating when the platform is moved and that hippocampal lesions disrupt this pattern^5,47^. In contrast, the present findings strongly and directly support the hypothesis that rearing is an ethological measure of hippocampal learning.

The current results also advance our understanding of the connections between rearing and hippocampal learning beyond work done on the topic by Wells et al.^18^ and Mun et al.^19^ since Lever and colleagues published their review. Wells et al. showed that increasing environmental novelty both decreases the speed modulation of hippocampal theta and increases the number of times rats reared, a result that indicated a coupling between hippocampal function and rearing behavior^18^. Mun et al.^19^ reported that the discrimination index in a novel place task (a measure of spatial memory) was positively related to rearing frequency during the initial exploration phase. Notably, this effect was specific to the novel place task, a task known to be sensitive to hippocampal integrity, and was not observed in the novel object task, a task that is generally insensitive to hippocampal integrity except at long delays^48,49^. Thus, rearing specifically promoted memory in a hippocampal dependent task. Neither Wells et al.^18^ nor Mun et al.^19^ tested the necessity of hippocampal activity during rearing for spatial memory. The results of our experiment are an advance over these works by showing explicitly that hippocampal activity during rearing is important for spatial memory.

While the function of rearing for spatial memory is not known, prior work suggests that the inactivation we performed in the current experiment may have disrupted spatial memory by interfering with updating of an internal model of the environment. Rearing is likely a form of active environmental sampling. As it relates to spatial memory, rearing has been suggested to aid in building and updating a model of the environment^15,24^. Rearing frequency is increased by cue changes that elicit hippocampal remapping^50^. Barth et al.^24^ suggested, based on analyses of hippocampal field potentials and unit activity, that the hippocampus switches to a distinct functional mode during rearing. This mode, they suggest, draws on sensory information gathered during the rearing to reduce uncertainty regarding allocentric position and to perform sensory realignment of the cognitive map^24^. By inactivating the hippocampus during rearing, our manipulation may have prevented this updating with downstream consequences for spatial memory. We note, however, that any updating is happening in our experiment is unlikely to be *de novo* environmental modeling. Prior to testing, while reaching performance criterion, our rats completed dozens of trials under identical circumstances providing ample time to model the environment proper. Importantly, however, the set of arms that opened during study (and which were still baited at test) varied randomly each day. Thus, the challenge on any given day was to disambiguate current trial information from the proactive interference of prior training. For this reason, we expect that any updating being performed would be to support trial specific event memory. This resembles the ‘Where did I park my car today?’ type memory, requiring disambiguation of recent events from similar prior events, that characterizes hippocampal function^51^.

The current work does not address if there is a specific portion of the hippocampus proper that is necessary for spatial memory encoding during rearing. However, prior work suggests the dentate gyrus may be important ^24,52,53^. Spatial memory ability in radial maze tasks is correlated to the size of mossy fiber terminal fields^52^. Separately, it was found that selectively breeding mice for frequent rearing behavior resulted in progeny with increased mossy fiber terminal fields^53^. The relevance of the dentate gyrus for driving hippocampal processing during rearing is also supported by the functional recordings^24^. Barth et al. ^24^ analyzed hippocampal current source density profiles during rearing and showed that rearing was accompanied by a prominent sink in the dentate gyrus and increased theta-gamma coupling in the terminal fields of the perforant path that originated in the medial entorhinal cortex^24^.

The present work also does not offer new insight into the dissociation between ‘supported’ and ‘unsupported’ rearing. ‘Supported’ and ‘unsupported’ refer to whether a rearing animal places its forelimbs on a surface while rearing to support itself. Prior work has shown these two forms of rearing to be dissociable. For example, selective breeding for unsupported rearing did not lead to a simultaneous increase in supported rearing^53^. Further, each supported and unsupported rearing are dissociable by their respective sensitivity to different motivating factors such as stress or motivation to escape^54,55^. The radial arm maze used in the present work, however, had narrow corridors and there were few occasions when the rats did not place a fore paw on a wall. Thus, though not formally analyzed, we casually observed that virtually all rearing events would have been classified as supported rearing. Moreover, the closed loop inactivation protocol did not dissociate between unsupported and supported in any case. Thus, separate work is required to determine if different results would have been obtained if inactivation would have been restricted to one type of rearing or the other.

Finally, regarding the specificity of our results, there are two sources of ambiguity worth noting. First, related to the discussion of supported versus unsupported rearing above, it may be that not every rearing event directly contributes to spatial memory encoding. Careful analysis of hippocampal dynamics across rearing events in future work could offer insight in this regard. Second, the current work does not address whether the specific form of encoding that occurs during rearing could happen in the absence of rearing. There are, for example, other behaviors that share a common phenotype with rearing that we call a ‘attentive sampling from place.’ For example, Monaco et al. described ‘lateral head-scanning’ behaviors that were coincident with endogenous formation of new place fields in hippocampal CA1 neurons^56^. In both rearing and lateral head-scanning, animals stand in one location and actively orient towards distal cues. Functionally, both are characterized by high amplitude theta in the hippocampus^24,56^. However, there is evidence to suggest that the encoding that occurs during rearing may not readily occur during non-rearing behaviors. Mun et al. induced inflammation in a mouse’s hind paw and found that it simultaneously hindered rearing and reduced sensitivity to spatial novelty^19^. Implicit to this result is that the mice did not switch to an alternate strategy to perform the same encoding when rearing was rendered uncomfortable. Thus, while the results of the present work are sufficient to demonstrate that rearing is an epoch of hippocampal dependent spatial memory encoding, additional work is needed to fully specify the relationship between rearing and spatial memory encoding.

Final summary and conclusion: This work demonstrates that disrupting activity in the dorsal hippocampus proper during rearing events is sufficient to disrupt spatial memory encoding in the delayed win-shift radial arm maze task. This result indicates that rearing events are an epoch of hippocampal dependent spatial memory encoding.

## Author Contributions

N.S., K.B., D.L. and E.N. designed the experiment. N.S., D.L., and K.B. collected all the data. D.L. and E.N. analyzed the data and wrote the manuscript.

## Acknowledgements

We would like to thank Charles Maitha for his technical support in setting up the 3D camera tracking system and laser control. We thank Indiana University Laboratory Animal Resources facilities for their attention and care of our animals. We thank Dr. Muriel Alejandra Mardones Diaz for advice and support on IHC. This work has been supported by the National Institutes of Health (by R01AG076198 to E.N), Indiana University Faculty Research Support Program (E.N.), Harlan Scholars Program (D.L.) and Hutton Honors College Research Grant (K.B).

## References

1. Olton, D. S., Walker, J. A., & Gage, F. H. (1978). Hippocampal connections and spatial discrimination. Brain research, 139(2), 295–308.

2. Jarrard, L. E. (1978). Selective hippocampal lesions: differential effects on performance by rats of a spatial task with preoperative versus postoperative training. journal of Comparative and Physiological Psychology, 92(6), 1119.

3. Olton, D. S. & Paras, B. C. Spatial memory and hippocampal function. Neuropsychologia, 17, 669–682 (1979).

4. Morris, R. G., Garrud, P., Rawlins, J. A., & O’Keefe, J. (1982). Place navigation impaired in rats with hippocampal lesions. Nature, 297(5868), 681–683.

5. Sutherland, R. J., Whishaw, I. Q. & Kolb, B. A behavioural analysis of spatial localization following electrolytic, kainate- or colchicine-induced damage to the hippocampal formation in the rat. Behav Brain Res, 7, 133–153 (1983).

6. Eichenbaum, H., Dudchenko, P., Wood, E., Shapiro, M. & Tanila, H. The Hippocampus, Memory, and Place Cells Is It Spatial Memory or a Memory Space? Neuron 23, 209–226 (1999).

7. Broadbent, N. J., Squire, L. R. & Clark, R. E. Spatial memory, recognition memory, and the hippocampus. Proc National Acad Sci 101, 14515–14520 (2004).

8. O’Keefe, J. & Dostrovsky, J. The hippocampus as a spatial map. Preliminary evidence from unit activity in the freely-moving rat. Brain Research, 34, 171–175 (1971).

9. O’Keefe, J. & Nadel, L. The Hippocampus as a Cognitive Map. (Oxford: Clarendon Press, 1978).

10. Dupret, D., O’Neill, J., Pleydell-Bouverie, B. & Csicsvari, J. The reorganization and reactivation of hippocampal maps predict spatial memory performance. Nature Neuroscience 13, 995–1002 (2010).

11. Foster, D. J. & Wilson, M. A. Reverse replay of behavioural sequences in hippocampal place cells during the awake state. Nature 440, 680–683 (2006).

12. Diba, K. & Buzsáki, G. Forward and reverse hippocampal place-cell sequences during ripples. Nature Neuroscience 10, 1241–1242 (2007).

13. Jadhav, S. P., Kemere, C., German, P. W. & Frank, L. M. Awake hippocampal sharp-wave ripples support spatial memory. Science (New York, NY) 336, 1454–1458 (2012).

14. Cheng, K. Oscillators and servomechanisms in orientation and navigation, and sometimes in cognition. Proc Royal Soc B Biological Sci 289, 20220237 (2022).

15. Lever, C., Burton, S. & O’Keefe, J. (2006) Rearing on Hind Legs, Environmental Novelty, and the Hippocampal Formation. Rev Neuroscience, 17, 111–134. doi: 10.1515/revneuro.2006.17.1-2.111

16. Thiel, C. M., Huston, J. P. & Schwarting, R. K. W. Hippocampal acetylcholine and habituation learning. Neuroscience 85, 1253–1262 (1998).

17. Anderson, M. I. et al. Behavioral correlates of the distributed coding of spatial context. Hippocampus 16, 730–742 (2006).

18. Wells, C.E., Amos, D.P., Jeewajee, A., Douchamps, V., Rodgers, J., O’Keefe, J., Burgess, N., and Lever, C. (2013). Novelty and anxiolytic drugs dissociate two components of hippocampal theta in behaving rats. J. Neurosci. 33, 8650–8667

19. Mun, H.S., Saab, B.J., Ng, E., McGirr, A., Lipina, T.V., Gondo, Y., Georgiou, J., and Roder, J.C. (2015). Self-directed exploration provides a Ncs1-dependent learning bonus. Sci. Rep.5, 17697.

20. Landfield, P. W., McGaugh, J. L., & Tusa, R. J. (1972). Theta rhythm: a temporal correlate of memory storage processes in the rat. Science, 175(4017), 87–89.

21. Robinson, T. E. & Vanderwolf, C. H. Electrical stimulation of the brain stem in freely moving rats: II. Effects on hippocampal and neocortical electrical activity, and relations to behavior. Exp Neurol 61, 485–515 (1978).

22. Berry, S. D., & Thompson, R. F. (1978). Prediction of learning rate from the hippocampal electroencephalogram. Science, 200(4347), 1298–1300.

23. Young, C. K. & McNaughton, N. Coupling of Theta Oscillations between Anterior and Posterior Midline Cortex and with the Hippocampus in Freely Behaving Rats. Cereb Cortex 19, 24–40 (2009).

24. Barth, A. M., Domonkos, A., Fernandez-Ruiz, A., Freund, T. F., & Varga, V. (2018). Hippocampal network dynamics during rearing episodes. Cell reports, 23(6), 1706–1715.

25. Hasselmo, M. E., Bodelón, C. & Wyble, B. P. A proposed function for hippocampal theta rhythm: separate phases of encoding and retrieval enhance reversal of prior learning. Neural Computation 14, 793–817 (2002).

26. Norman, K. A., Newman, E. L. & Detre, G. A neural network model of retrieval-induced forgetting. 114, 887–953 (2007).

27. Montgomery, S. M., Betancur, M. I., & Buzsáki, G. (2009). Behavior-dependent coordination of multiple theta dipoles in the hippocampus. Journal of Neuroscience, 29(5), 1381–1394.

28. Richard, G. R., Titiz, A., Tyler, A., Holmes, G. L., Scott, R. C., & Lenck-Santini, P. P. (2013). Speed modulation of hippocampal theta frequency correlates with spatial memory performance. Hippocampus, 23(12), 1269–1279.

29. Schmidt, B., Hinman, J. R., Jacobson, T. K., Szkudlarek, E., Argraves, M., Escabí, M. A., & Markus, E. J. (2013). Dissociation between dorsal and ventral hippocampal theta oscillations during decision-making. Journal of Neuroscience, 33(14), 6212–6224.

30. Douchamps, V., Jeewajee, A., Blundell, P., Burgess, N., & Lever, C. (2013). Evidence for encoding versus retrieval scheduling in the hippocampus by theta phase and acetylcholine. Journal of Neuroscience, 33(20), 8689–8704.

31. Belchior, H., Lopes-dos-Santos, V., Tort, A. B., & Ribeiro, S. (2014). Increase in hippocampal theta oscillations during spatial decision making. Hippocampus, 24(6), 693–702.

32. Siegle, J. H., & Wilson, M. A. (2014). Enhancement of encoding and retrieval functions through theta phase-specific manipulation of hippocampus. elife, 3.

33. Hernández-Pérez, J. Jesús, Blanca E. Gutiérrez-Guzmán, and Maria, E. Olvera-Cortés. “Hippocampal strata theta oscillations change their frequency and coupling during spatial learning.” Neuroscience 337 (2016): 224–241.

34. Honey, C. J., Newman, E. L. & Schapiro, A. C. Switching between internal and external modes: A multiscale learning principle. Network Neuroscience 1, 339–356 (2017).

35. Winson, J. (1978). Loss of hippocampal theta rhythm results in spatial memory deficit in the rat. Science, 201(4351), 160–163.

36. Mitchell, S. J., Rawlins, J. N., Steward, O., & Olton, D. S. (1982). Medial septal area lesions disrupt theta rhythm and cholinergic staining in medial entorhinal cortex and produce impaired radial arm maze behavior in rats. Journal of Neuroscience, 2(3), 292–302.

37. Mizumori, S. J. Y., Perez, G. M., Alvarado, M. C., Barnes, C. A., & McNaughton, B. L. (1990). Reversible inactivation of the medial septum differentially affects two forms of learning in rats. Brain research, 528(1), 12–20.

38. Wang, Y., Romani, S., Lustig, B., Leonardo, A., & Pastalkova, E. (2015). Theta sequences are essential for internally generated hippocampal firing fields. Nature neuroscience, 18(2), 282–288.

39. McNaughton, N., Ruan, M., & Woodnorth, M. A. (2006). Restoring theta-like rhythmicity in rats restores initial learning in the Morris water maze. Hippocampus, 16(12), 1102–1110.

40. Shirvalkar, P. R., Rapp, P. R., & Shapiro, M. L. (2010). Bidirectional changes to hippocampal theta-gamma comodulation predict memory for recent spatial episodes. Proceedings of the National Academy of Sciences, 107(15), 7054–7059.

41. Packard, M. G., Regenold, W., Quirion, R., & White, N. M. (1990). Post-training injection of the acetylcholine M 2 receptor antagonist AF-DX116 improves memory. Brain Research, 524(1), 72–76

42. Seamans, J. K., & Phillips, A. G. (1994). Selective memory impairments produced by transient lidocaine-induced lesions of the nucleus accumbens in rats. Behavioral neuroscience, 108(3), 456.

43. Gradinaru V, Zhang F, Ramakrishnan C, Mattis J, Prakash R, Diester I, Goshen I, Thompson KR, Deisseroth K. Molecular and cellular approaches for diversifying and extending optogenetics. Cell. 2010;141(1):154–65.

44. Weitz, A. J., Fang, Z., Lee, H. J., Fisher, R. S., Smith, W. C., Choy, M., … & Lee, J. H. (2015). Optogenetic fMRI reveals distinct, frequency-dependent networks recruited by dorsal and intermediate hippocampus stimulations. Neuroimage, 107, 229–241.

45. Deacon, R. M. J., Croucher, A. & Rawlins, J. N. P. Hippocampal cytotoxic lesion effects on species-typical behaviours in mice. Behav Brain Res 132, 203–213 (2002).

46. Kamei, C., Chen, Z., Nakamura, S. & Sugimoto, Y. Effects of intracerebroventricular injection of histamine on memory deficits induced by hippocampal lesions in rats. Method Find Exp Clin, 19, 253–9 (1997).

47. Sutherland, R. J. R., Whishaw, I. Q. I. & Regehr, J. C. J. Cholinergic receptor blockade impairs spatial localization by use of distal cues in the rat. Journal of comparative and physiological psychology, 96, 563–573 (1982).

48. Broadbent NJ, Gaskin S, Squire LR, Clark RE. Object recognition memory and the rodent hippocampus. Learn Mem. 2009 Dec 22;17(1):5–11

49. Cohen SJ, Stackman RW Jr. Assessing rodent hippocampal involvement in the novel object recognition task. A review. Behav Brain Res. 2015 May 15;285:105–17.

50. Wells, C. E., Krikke, B., Saunders, J., Whittington, A. & Lever, C. Changes to open field surfaces typically used to elicit hippocampal remapping elicit graded exploratory responses. Behav Brain Res 197, 234–238 (2009).

51. Knierim, J. J., Lee, I., & Hargreaves, E. L. (2006). Hippocampal place cells: parallel input streams, subregional processing, and implications for episodic memory. Hippocampus, 16(9), 755–764.

52. Jamot L, Bertholet JY, Crusio WE. Neuroanatomical divergence between two substrains of C57BL/6J inbred mice entails differential radial-maze learning. Brain Res 1994; 644: 352–356.

53. Crusio WE. Genetic dissection of mouse exploratory behaviour. Behav Brain Res 2001; 125: 127–132.

54. Sturman, O., Germain, P. L., & Bohacek, J. (2018). Exploratory rearing: a context-and stress-sensitive behavior recorded in the open-field test. Stress, 21(5), 443–452.

55. Griebel G, Blanchard DC, Blanchard RJ. Evidence that the behaviors in the Mouse Defense Test Battery relate to different emotional states: a factor analytic study. Physiol Behav 1996; 60: 1255–1260.

56. Monaco, J. D., Rao, G., Roth, E. D., & Knierim, J. J. (2014). Attentive scanning behavior drives one-trial potentiation of hippocampal place fields. Nature Neuroscience, 17(5), 725–731. https://doi.org/10.1038/nn.3687

